# A Bayesian approach to Mendelian randomization using summary statistics in the univariable and multivariable settings with correlated pleiotropy

**DOI:** 10.1101/2023.05.30.542988

**Authors:** Andrew J. Grant, Stephen Burgess

## Abstract

Mendelian randomization uses genetic variants as instrumental variables to make causal inferences on the effect of an exposure on an outcome. Due to the recent abundance of high-powered genome-wide association studies, many putative causal exposures of interest have large numbers of independent genetic variants with which they associate, each representing a potential instrument for use in a Mendelian randomization analysis. Such polygenic analyses increase the power of the study design to detect causal effects, however they also increase the potential for bias due to instrument invalidity. Recent attention has been given to dealing with bias caused by correlated pleiotropy, which results from violation of the Instrument Strength independent of Direct Effect assumption. Although methods have been proposed which can account for this bias, a number of restrictive conditions remain in many commonly used techniques. In this paper, we propose a novel Bayesian framework for Mendelian randomization which provides valid causal inference under very general settings. We propose the methods MR-Horse and MVMR-Horse, which can be performed without access to individual-level data, using only summary statistics of the type commonly published by genome-wide association studies, and can account for both correlated and uncorrelated pleiotropy. In simulation studies, we show that the approach retains type I error rates below nominal levels even in high pleiotropy scenarios. We consider an applied example looking at the causal relationship between combinations of four exposures (LDL-cholesterol, triglycerides, fasting glucose and birth weight) and three outcomes (coronary artery disease, type 2 diabetes and asthma).

## Introduction

Mendelian randomization is a technique which exploits genetic variation to make causal inferences on the effect of one or more exposures on an outcome of interest (Lawlor et al., 2008). The underlying idea is that, because of the Mendelian laws of inheritence, a genetic variant which is associated with an exposure of interest can serve as a proxy for the exposure. Such a proxy should be less susceptible to the counfounding and reverse causation which can inhibit causal inference from observational data. Methods for Mendelian randomization have been developed which can combine information from multiple genetic variants associated with the same exposure. Furthermore, many of these methods can be performed using only summary statistics taken from genome-wide association studies (GWAS), removing the need to access sensitive individual level data. Under the two-sample framework, summary statistics for the exposure and outcome can be taken from separate, ideally non-overlapping, samples (Pierce and Burgess, 2013). Thus, any combination of exposure and outcome for which there exists publicly available GWAS summary statistics can potentially be considered in a Mendelian randomization study. However, in order to provide robust evidence for causal effects, strong assumptions must be made pertaining to the relationships between the genetic variants and the exposures, outcome and any confounder variables.

Mendelian randomization can be considered as a form of instrumental variables analysis, where each genetic variant must satisfy the three instrumental variables assumptions. In the single exposure case, these assumptions are that each genetic variant:

1. is associated with the exposure;
2. is not associated with the outcome via any confounding pathway;
3. does not affect the outcome other than via the exposure.

The relationships between a valid genetic variant and other relevant traits is illustrated graphically in the directed acyclic graph (DAG) in Figure 1A (Greenland, 2000).

**Figure 1:**
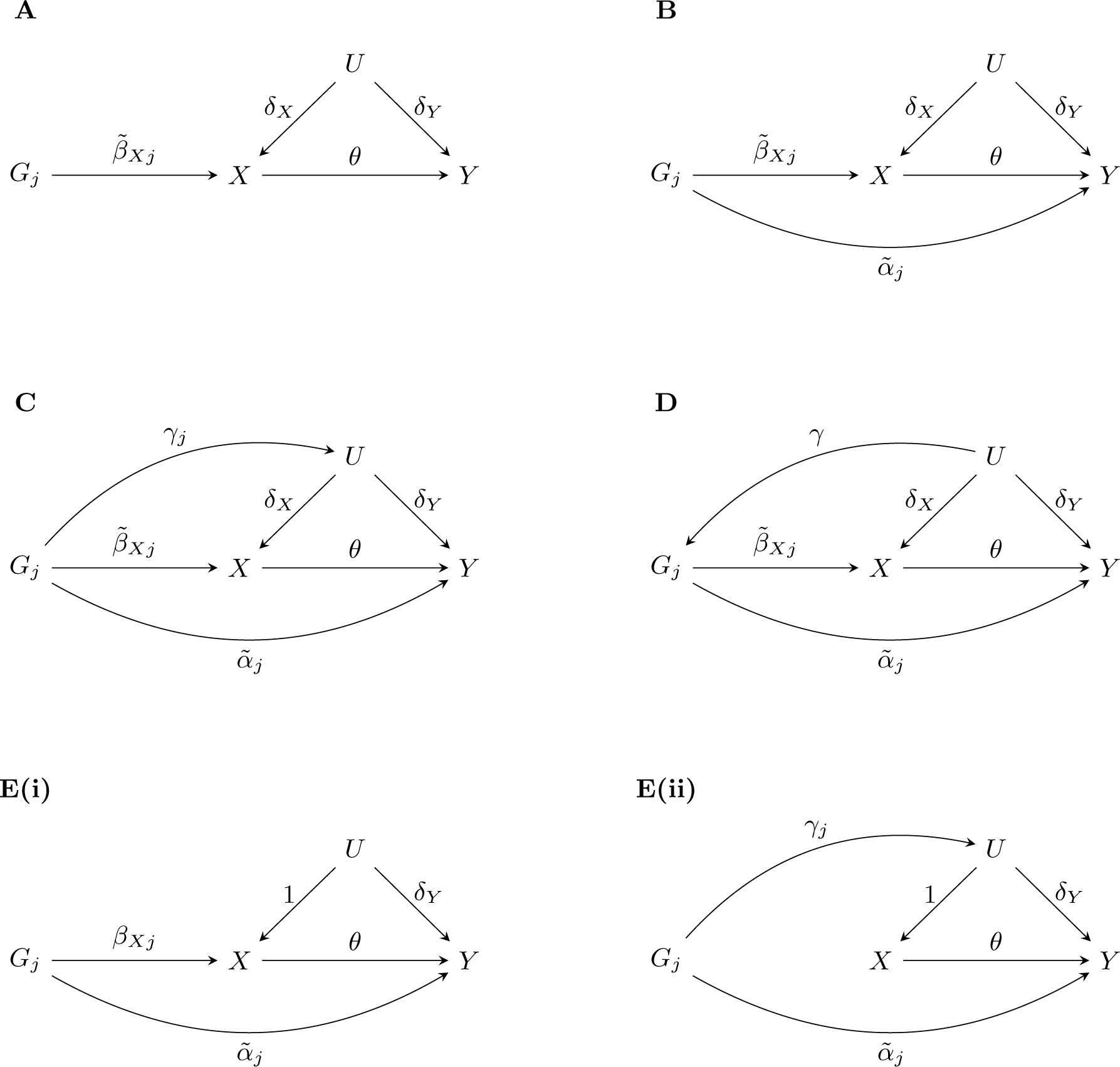
Directed acyclic graphs showing the relationship between the *j*th genetic variant (*G_j_*), exposure (*X*), outcome (*Y*) and unmeasured confounders (*U*) when: (A) *G_j_* is a valid instrument; (B) *G_j_* is invalid due to a direct effect on *Y* ; (C) *G_j_* is invalid due to both a direct effect on *Y* and an indirect effect via *U* ; (D) *G_j_* is invalid due to both a direct effect on *Y* and being caused by *U* ; (E) *G_j_* is invalid due to either (i) a direct effect on *Y* or (ii) both a direct effect on *Y* and an indirect effect via *U* but with no direct effect on *X*.

A common source of instrument invalidity is pleiotropy, which is where genetic variants associate with multiple traits. If a genetic instrument associates with a predictor of the outcome via a causal pathway which bypasses the exposure, then this will open up a pleiotropic pathway and invalidate the third instrumental variables assumption. Another source of invalidity is population structure, which is where there are differences in the genetic distributions among subgroups of the population being studied. This results in confounding of the genetic variant-outcome relationship, and invalidates the second instrumental variables assumption. Population structure may be caused by, for example, population stratification, dynastic effects and assortative mating (Brumpton et al., 2020).

Much attention has been given in the literature to developing methods for Mendelian randomization which are robust to instrument invalidity. Many of these methods have been developed motivated by providing robustness against pleiotropy. Often, the InSIDE (Instrument Strength Independent of Direct Effects) assumption is required for these methods to give valid results, which means that any pleiotropic effects are independent of the genetic variant-exposure associations (Bowden et al., 2015). Methods which require InSIDE will not provide robustness to population structure, since the confounding pathway may create a correlation between the genetic effects on the exposure and the direct effects on the outcome. Recently, more attention has been given to methods which can also handle violations of the InSIDE assumption, a scenario referred to as correlated pleiotropy (Morrison et al., 2020). However, methods which provide such robustness typically suffer from a combination of inflated type I error rates, low power or further restrictive assumptions.

In this paper we propose a novel Bayesian approach to Mendelian randomization which is robust to both uncorrelated and correlated pleiotropy. We show that this approach is able to provide unbiased and efficient causal effect estimation without inflated type I error rates in very general settings, with favorable performance in comparison with existing methods. A further advantage of the proposed method is that it is easily generalised to the multivariable Mendelian randomization paradigm, which has far fewer robust methods than the single exposure case. We demonstrate the approach in an applied example which looks at the relationship between combinations of four exposures (LDL-cholesterol, triglycerides, fasting glucose and birth weight) and three outcomes (coronary artery disease, type 2 diabetes and asthma).

## Methods

### Models for instrument invalidity

We consider a potentially casual exposure *X* for an outcome *Y* with unmeasured confounders *U* . We denote by *θ* the causal effect of *X* on *Y*, and assume that this effect is linear and homogeneous in the population (that is, that there is no effect modification). This latter assumption allows for causal effect estimation in the instrumental variables framework.

The relationships between a valid genetic instrument, *G_j_*, and the exposure, outcome, and unmeasured confounders, are shown in Figure 1A. If we have a single such instrument, we can estimate the causal effect using the Wald ratio, 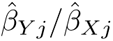, where 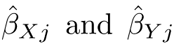 are estimates of the associations between *G_j_* and *X* and *Y*, respectively. These association estimates are typically obtained by using linear regression for continuous traits and logistic regression for binary traits. If we have *J* valid instruments, the inverse-variance weighted (IVW) method fits the model

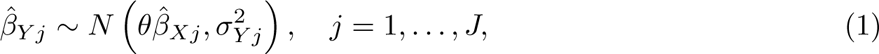

where *σ_Y j_* is the standard error of the *j*th genetic variant-outcome association estimate (Burgess et al., 2013). Note that the standard error of the *j*th genetic variant-exposure association estimate, *σ_Xj_*, does not appear in the IVW model, which reflects the fact that this approach implicitly assumes that the genetic associations with the exposure are measured without error.

Most pleiotropy robust methods for Mendelian randomization make the assumption that some of the instruments are valid, and allow for varying levels of invalidity. MR-PRESSO assumes that a small number of instruments are invalid which will result in outliers in the Wald ratio estimates (Verbanck et al., 2018). Under the InSIDE assumption, MR-PRESSO tests for outliers and estimates the causal effect using IVW with the identified outliers removed. If at least half of the instruments are valid, a weighted median of the Wald ratios gives a consistent estimate of the causal effect, whether or not InSIDE is satisfied (Bowden et al., 2016).

Figure 1B illustrates the case where there is a pleiotropic pathway to the outcome which does not pass via *X* or *U* (although it may go via other unmeasured traits which do not associate with the exposure). In this case, the direct effect *α*~*_j_* is independent of the instrument strength 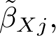 hence InSIDE is met. The causal effect can be estimated using the MR-Egger method, which fits the model

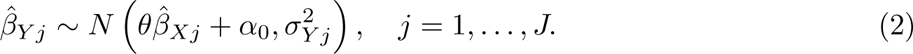

The intercept term *α*_0_ represents the average pleiotropic effect across all variants. If this average effect is equal to zero, then it is said that pleiotropy is balanced. If the average effect is non-zero, the pleiotropy is said to be directional. Under the MR-Egger model, potentially all variants may be invalid, but only in such a way that the InSIDE assumption is satisfied.

Figure 1C shows the case where the genetic variant has a pathway to the outcome via one or both of a direct path and a path that passes through a confounder of the exposure-outcome relationship. This latter pathway creates correlated pleiotropy, or, in other words, violates InSIDE. One commonly used model is for this scenario is

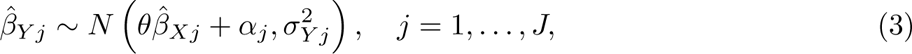

where *α_j_* represents the effect 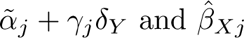 and 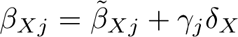. Under the assumption that some of the instruments are valid and some are invalid, Kang et al. (2016) and Rees et al. (2019) proposed estimating *θ* using penalised regresssion with a lasso-style penalty on the *α_j_* terms. In this way, the estimates of *α_j_* relating to the valid instruments are shrunk toward zero. A post-lasso approach fits the IVW model using only the instruments whose *α_j_* estimates were shrunk to zero in the initial lasso regression. Although these lasso-based approaches provide unbiased effect estimates, they can suffer from substantially inflated type I error rates due to the post-selection inference problem.

Kang et al. (2016) showed that the causal effect is identifiable in the model given in equation (3) when a plurality of instruments are valid. That is, if there are groups of variants for which the single-instrument causal effect estimate is the same, the largest of these groups is the set of valid instruments. Alternative methods that make the plurality valid assumption include the Contamination Mixture method (Burgess et al., 2020) and MR-Mix (Qi and Chatterjee, 2019), which are based on fitting mixture distributions to the Wald ratio estimates, and weighted mode approaches (Hartwig et al., 2017; Guo et al., 2018).

Xue et al. (2021) proposed a method which fits the model

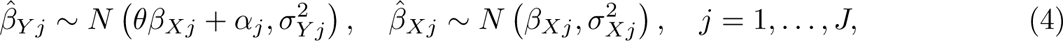

using constrained maximum likelihood, referred to as MR-cML. This method provides consistent causal effect estimation under the plurality valid assumption. Similar to the lasso regression approaches, MR-cML maximises the likelihood in such a way that the *α_j_*’s pertaining to valid instruments will tend toward zero. However, through the use of model averaging it suffers less from inflated type I errors. Furthermore, because the model incorporates the *σ_Xj_* terms, it provides robustness to weak instruments. The MR-RAPS method is also based on the model given in equation (4), where pleiotropy is accounted for using a robust loss function. However, MR-RAPS makes the InSIDE assumption, requires balanced pleiotropy, and was shown to have lower power than MR-cML by Xue et al. (2021).

Bayesian methods have been far less utilised in the Mendelian randomization literature, despite a number of potential advantages over frequentist approaches in fitting a model such as that given in equation (4). Berzuini et al. (2020) developed a Bayesian approach where parameters representing pleiotropic effects are given a shrinkage prior. This has the effect of shrinking these terms pertaining to the valid instruments toward zero, similar to the lasso-based approaches. The use of credible intervals obtained from the posterior distribution was shown to provide better coverage than the weighted median method, suggesting that the Bayesian approach is better able to estimate the uncertainty in the effect estimates. The method requires individual-level data and an assumption akin to InSIDE. Zhao et al. (2019) (BWMR) and Bucur et al. (2020) (BAYESMR) have proposed Bayesian approaches using summary-level data. Similar to Berzuini et al. (2020), these approaches place a shrinkage prior on the *α_j_* terms and make the InSIDE assumption.

Recently developed approaches aimed at the correlated pleiotropy case include CAUSE (Morrison et al., 2020), MR-Corr2 (Cheng et al., 2021), and MR-CUE (Cheng et al., 2022), which each utilise Bayesian methods. MR-Corr2 uses a spike-and-slab prior for pleiotropic terms, similar to BAYESMR, and also includes a parameter to account for correlated pleiotropy. However, it does not allow for both uncorrelated and correlated pleiotropy among the variants. CAUSE and MR-CUE make the assumption, illustrated in Figure 1E, that genetic instruments are either directly associated with the exposure or only associated with the exposure via a confounder. Xue et al. (2021) showed that CAUSE is particularly sensitive to violations of this assumption. Another feature of these approaches is that they utilise genome-wide summary statistics, as opposed to just’summary statistics relating to a curated set of instruments. The genome-wide summary statistics are used to estimate nuisance parameters which are inputs to the Bayesian models.

Figure 1D shows the case where correlated pleiotropy is caused by a confounder of the genetic variant-outcome relationship, for example due to population structure. Although the cause of instrument invalidity differs to that shown in Figure 1C, we can model it in the same way. We thus consider (4) to be a very general parametric model for Mendelian randomization with pleiotropy. It can represent invalidity due to both pleiotropy and population structure, and does not make the assumption of CAUSE and MR-CUE that the instruments cannot both be directly associated with the exposure and subject to correlated pleiotropy.

### Univariable Mendelian randomization

Suppose we have, for *J* genetic variants, summary statistics of their association with the exposure and outcome. We denote the estimate of the association between the *j*th variant and the exposure and outcome by 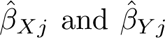, respectively, and the standard errors of the estimates by *σ_Xj_* and *σ_Y j_*. We model the association estimates by

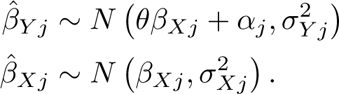

We place a jointly normal prior on (*β_Xj_, α_j_*)

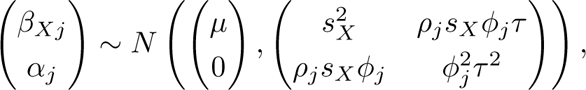

and we use the following weakly informative priors for the other unknown parameters

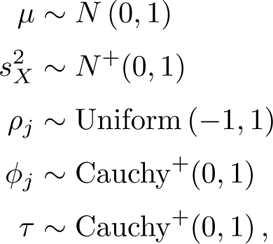

where *N* ^+^(*·, ·*) and Cauchy^+^(*·, ·*) are the half-normal and half-Cauchy distributions, respectively. We also place a uniform prior on *θ* with wide limits, in our case *±*10.

In this model, the *β_Xj_* term represents the effect of the *j*th genetic variant on the exposure, and the *α_j_* term represents the effect of the variant on the outcome which bypasses the exposure. If the *j*th variant is a valid instrument, then the true value of *α_j_* is zero. Otherwise, the variant is pleiotropic, and the pleiotropic effect is accounted for by a non-zero *α_j_*. If the *j*th variant is either a cause of, or caused by, *U*, then *β_Xj_* and *α_j_* will be correlated. This correlation is modelled by the *ρ_j_* parameter. Thus, the model can account for valid instruments (*α_j_* = 0), uncorrelated pleiotropy (*α_j_* ≄ 0*, ρ_j_* = 0) and correlated pleiotropy (*α_j_* ≄ 0*, ρ* ≄ 0).

Under this parametrisation, the marginal distribution of *α_j_* is the horseshoe prior (Carvalho et al., 2010). The horseshoe is a shrinkage prior which will shrink some of the *α_j_*’s toward zero, while non-zero *α_j_*’s will escape shrinkage better than when using a Laplace prior (which is equivalent to lasso regression). Thus, it is less likely to underestimate the value of non-zero *α_j_*’s. The *τ* parameter controls the global level of shrinkage such that a lower value will shrink more of the *α_j_*’s close to zero. In our case, we assign *τ* a half-Cauchy prior and allow its posterior distribution to be estimated from data. However, if there was prior knowledge on the level of sparsity, this could be incorporated by adjusting this prior or setting a fixed value (Piironen and Vehtari, 2017). The horsehoe prior has previously been proposed for pleiotropic effects in Mendelian randomization by Berzuini et al. (2020) for the individual level data setting.

In practice, we have implemented the method using BUGS via the rjags package in R (Plummer, 2022). We take 10 000 samples from the posterior distribution (with a burn-in of 10 000 iterations) from 3 chains, which, in all cases reported in the simulation study and applied examples, resulted in R-hat values for the parameters of interest close to 1, indicating good convergence. The causal effect estimate is the mean of the posterior samples, and the 95% credible interval is the lower and upper 2.5th percentiles. We call our method MR-Horse (after the *horse*shoe prior).

### Multivariable Mendelian randomization

Multivariable Mendelian randomization considers the effect of multiple exposures on the outcome (Burgess and Thompson, 2015; Burgess et al., 2015). Originally motivated as a method for accounting for measured pleiotropy, it can also be used to simultaneously estimate causal effects of multiple correlated exposures (Sanderson et al., 2019), to perform risk factor selection (Zuber et al., 2020) and in mediation analysis (Carter et al., 2021). A genetic variant is a valid instrumental variable for multivariable Mendelian randomization if: it is associated with at least one exposure; it is not associated with the outcome via any confounding pathway; and it does not affect the outcome other than via one or more of the measured exposures.

The multivariable IVW approach fits the model given in equation (1), but where the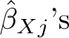 are vectors of the association estimates between the *j*th genetic variant and each exposure. There are far fewer pleiotropy robust methods for multivariable Mendelian randomization compared with the univariable setting. Commonly used approaches include multivariable versions of MR-Egger (Rees et al., 2017) and MR-PRESSO, a multivariable median-based approach (Grant and Burgess, 2021), and GRAPPLE (Wang et al., 2021), akin to a multivariable version of MR-RAPS. Recently, a multivariable version of MR-cML has been developed (Lin et al., 2023).

Our proposed model for univariable Mendelian randomization can be easily generalised to the multivariable case as follows. We denote by *β_Xj·_* the vector of associations between the *j*th genetic variant and the *K* exposures, and by 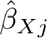 heir estimates which have covariance matrix Σ*_Xj_*. As with the standard errors in the univariable case, the Σ*_Xj_*’s are assumed known. In practice, they can be estimated either using individual level data or from genome-wide summary statistics (see, for example, Sanderson et al., 2021 and Grant et al., 2022). We fit the model

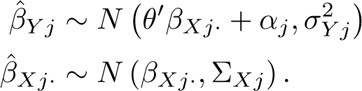

We place a joint prior on the (*β_Xj·_, α_j_*) as follows

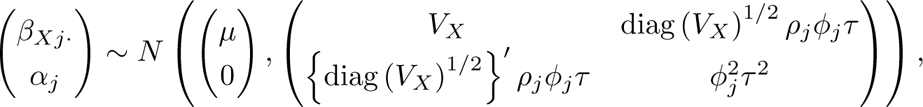

and we use the following non-informative priors for the other unknown parameters

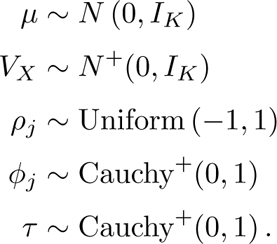

As in the univariable case, this parametrisation puts a horseshoe shrinkage prior on the *α_j_*’s with global shrinkage controlled by the *τ* parameter. The correlation between the genetic variant-exposure effects and the pleiotropic effects are accounted for via the *ρ_j_* parameters.

As before, we have implemented the method using BUGS and the rjags package in R, with 10 000 samples taken from the posterior distribution (with a burn-in of 10 000 iterations) from 3 chains. In all cases reported in the simulation study, R-hat values for the parameters of interest were close to 1, indicating good convergence. We call the multivariable method MVMR-Horse.

### Simulation study

#### Primary simulation parametrisation

In order to examine the performance of our method in comparison with alternatives, we generated data based on the simulation parametrisation of Xue et al. (2021), with some parameters adapted to ensure levels of instrument strength and variation of the exposure explained that is typical of realistic settings. The data generating mechanism is

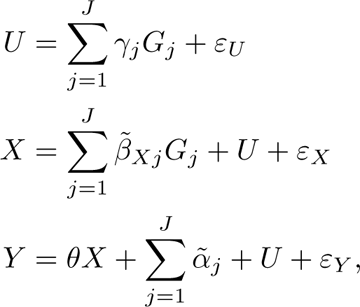

where *G_j_ ∼* Binomial (2*, f_j_*), *f_j_ ∼* Uniform (0.1, 0.3), and *ε_U_, ε_X_, ε_Y_ ∼ N* (0, 1) independently. We considered scenarios with *J* = 20 or 100 genetic variants. For each *j*, 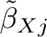 was generated uniformly on the interval (*−*0.15, − 0.05) *∪* (0.05, 0.15), *γ_j_ ∼* Uniform (*−*0.1, 0.1) and *δ_X_* = *δ_Y_* = 1. We considered scenarios where either 20%, 40% or 60% of variants were invalid instruments. Unlike Xue et al. (2021), we considered both balanced and directional pleiotropy scenarios. In the balanced pleiotropy case, *α_j_ ∼* Uniform (*−*0.2, 0.2), and in the directional pleiotropy case, *α_j_ ∼* Uniform (*−*0.1, 0.3), when the *j*th variant was invalid, and in both cases *α_j_* = 0 when the *j*th variant was valid. The causal effect was either *θ* = 0.1 or 0. Across the *J* = 100 scenarios, the average *R*^2^ value was between 15.4% and 16.9%, and the mean F-statistic ranged between 77.4 and 84.9. Across the *J* = 20 scenarios, the average *R*^2^ value was between 3.5% and 3.9%, and the mean F-statistic ranged between 88.0 and 98.2. Note that under this parametrisation, InSIDE was violated in all scenarios.

In each of 1 000 replications, two independent samples of size *n* = 50 000 were generated. The genetic association estimates with the exposure were taken in one sample, and with the outcome in the other. The association estimates and their standard errors were obtained using simple linear regression of the exposure and outcome on each genetic variant in turn.

#### Comparison methods

We compared our approach with standard IVW estimation and also an oracle estimator which performed IVW using only the truly valid instruments. We used a weighted median method, MR-Median, which requires half of the variants to be valid but does not make the InSIDE assumption nor requires balanced pleiotropy. We used BWMR to compare with another Bayesian approach, noting that this method assumes InSIDE. Finally, we compared with MR-cML as well as an extension to this approach MR-cML-DP. These latter two methods both make the plurality valid assumption and aim to shrink estimates of the pleiotropic effects pertaining to valid instruments toward zero. The extension uses data perturbation (DP) to produce multiple perturbed samples (the default, which we use, being 200 samples), and then takes averages across the samples for estimation of the causal effect and standard error. This significantly increases the computational time, but has been shown to reduce bias and type I error rates.

Other common methods not considered include MR-Egger, MR-RAPS and MR-PRESSO (InSIDE and balanced pleiotropy) and also ConMix, MR-Mix, MR-Mode, MR-Lasso (plurality valid, can handle some directional pleiotropy). In the simulation study of Xue et al. (2021), MR-cML generally outperformed each of these methods in terms of bias, power and type I error rates. Because our simulation study is largely based on that of Xue et al. (2021), it is expected that MR-cML would be the best performing out of these, and thus for brevity we don’t compare with all these methods in our study. Note that MR-cML also performed favorably compared with MR-Median in the study of Xue et al. (2021), however we retain MR-Median here so we have another comparison method which does not require balanced pleiotropy or InSIDE.

#### Genome-wide simulation study

We performed a simulation study which simulates a set of summary statistics representing genetic associations from independent variants across the genome, in order to compare our approach with CAUSE. We simulated *J* = 100 000 genetic variant associations such that either *p* = 100 or 20 variants were expected to have non-zero association with the outcome, setting the heritabilities of *X* and *Y* to *h_X_* = *h_Y_* = 0.3, respectively. We considered scenarios where either 20%, 40%, or 60% of these variants were expected to be pleiotropic. To do this, we first simulated 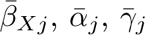 from the joint mixture distribution

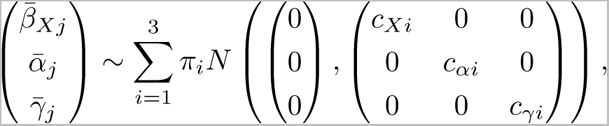

Where

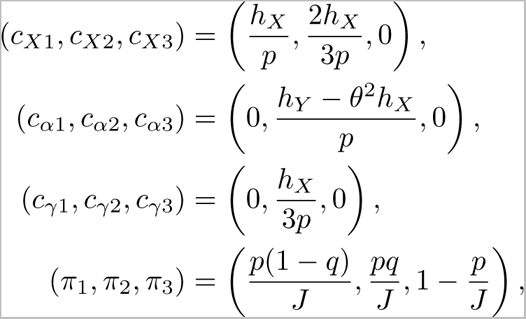

and *q* is the proportion of variants associated with the exposure which are invalid. These were converted to non-standardised effect sizes by 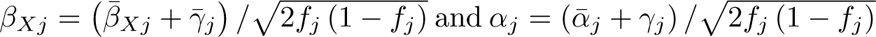.

The genetic association estimates were then generated by 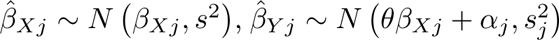, where 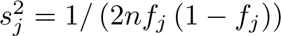. We considered scenarios where *θ* = 0.1 or 0.

For each variant, we computed a p-value using 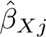 and *s_j_*. We performed the same methods used in the primary simulation, taking as inputs variants with p-value *<* 5 *×* 10*^−^*^8^ (that is, which pass the conventional genome-wide significance threshold). We also performed CAUSE using all simulated genetic variants, and with its default threshold p-value *<* 0.001 as classifying that a variant is associated with the exposure. Due to the extra computational time required to run the simulations, we restricted this simulation study to 200 replications per scenario.

#### Multivariable simulation study

To compare methods in the multivariable setting, we simulated from the same model as in the primary simulation with *J* = 100, but where there were two exposures generated according to

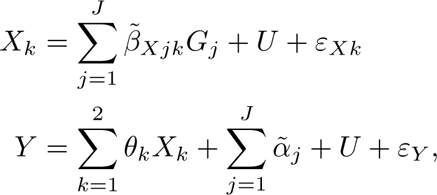

for *k* = 1, 2, where *θ*_1_ = 0.1 or 0, *θ*_2_ = 0.1, and (*ε_Xk_, ε_Xk_*) were simulated from the bivariate normal distribution with marginal variances of 1 and covariance of 0.3.

As well as the multivariable IVW and multivariable IVW-oracle methods, we also compared with the multivariable median approach (MVMR-Median), GRAPPLE and the constrained maximum likelihood approach with data perturbation (with 100 perturbed samples), which we denote MVMR-cML-DP. Note that for the methods which require the full covariance matrices of the genetic variant-exposures association estimates (that is, GRAPPLE, MVMR-cML-DP and MVMR-Horse), we input these assuming that the traits are independent, even though they are not. This it to reflect the typical scenario where only GWAS summary statistics are available without trait correlation estimates. Due to the extra computational time required to run the simulation, we restricted this simulation study to 200 replications per scenario.

### Applied example

We considered combinations of four exposures: LDL-cholesterol (LDL), triglycerides (TG), fasting glucose (FG); and birth weight (BW), on three outcomes: coronary artery disease (CAD); type 2 diabetes (T2D); and Asthma. Each of these exposure-outcome relationships were considered by both Morrison et al. (2020) and Xue et al. (2021), some with conflicting results. Following Morrison et al. (2020), we classify the exposure-outcome pairs as either: having a causal effect supported by the literature (LDL-CAD, TG-CAD, FG-T2D); having unknown or conflicting evidence of a causal effect (BW-CAD, TG-T2D, FG-CAD, BW-T2D, LDL-T2D); or having implausible, or unsupported evidence of a, causal effect (LDL-Asthma, FG-Asthma, BW-Asthma, TG-Asthma).

We analysed each of the exposure-outcome pairs using two-sample Mendelian randomization from publicly available GWAS results, as shown in Table 5. All genetic associations were estimated in samples of individuals of European, or predominantly European, ancestry. Note that some of these GWAS datasets differ to those used in Morrison et al. (2020) and Xue et al. (2021), and so there are some differences in the results that we report here.

**Table 5:**
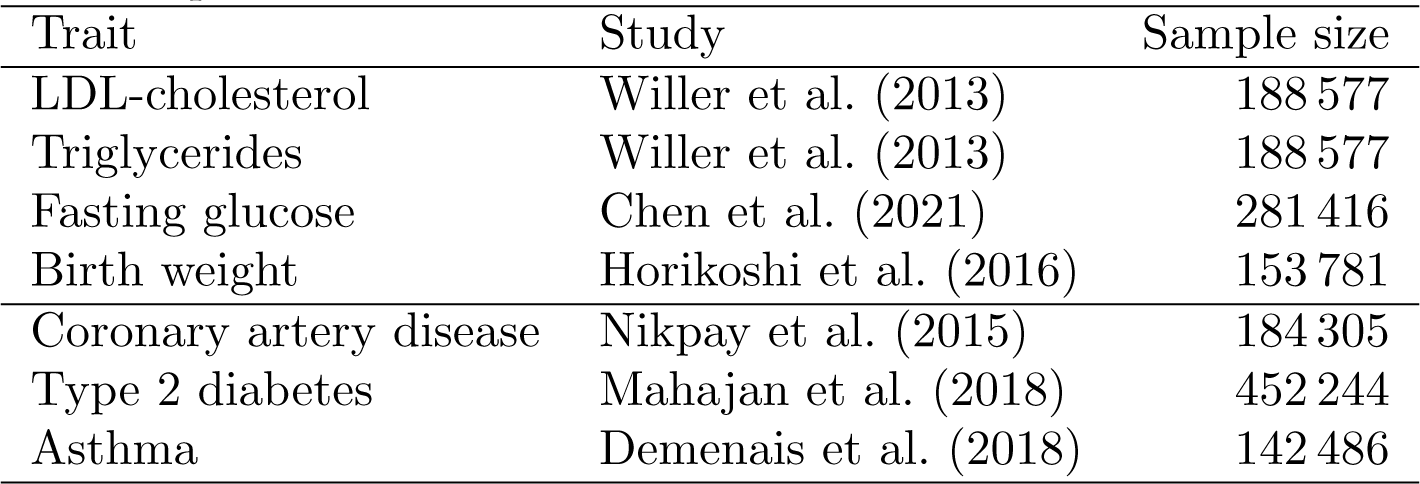
Sources of genetic association estimates for the four exposures and three outcomes considered in the applied example.

For each exposure, we chose a set of instruments by first selecting all genetic variants which associated with the trait with a p-value *<* 5 *×* 10*^−^*^8^, then pruned these to have *r*^2^ *<* 0.001. The set of variants was then harmonised with each outcome dataset, and only variants which appeared in both exposure and outcome datasets were retained. We performed analysis using the IVW, CAUSE, MR-cML, MR-cML-DP, and MR-Horse methods. We used the TwoSampleMR R package (Hemani et al., 2018) for the pruning and harmonisation procedures, and the MendelianRandomization R package (Yavorska and Burgess, 2017; Broadbent et al., 2020) for implementing the IVW, MR-cML and MR-cML-DP methods.

## Results

### Simulation study results

The mean estimates, standard deviation of estimates, coverage and power / type I error rates for each scenario are shown in Table 1 (*J* = 100), Table 2 (*J* = 20), Table 3 (genome-wide study) and Table 4 (multivariable setting, showing results relating to estimates of *θ*_1_ only). For MR-Horse and MVMR-Horse, we computed coverage as the proportion of replications for which the true causal effect was within the 95% credible interval, and rejection rate as the proportion of replications for which the 95% credible interval did not contain zero. For the other methods, except for CAUSE, we used the confidence intervals (or credible intervals) produced by their respective software packages to evaluate coverage and rejection rates in the same way (where 95% confidence intervals for BWMR were produced using the effect estimates and standard errors given by the BWMR package). For CAUSE, we evaluated coverage using the 95% credible interval, and the rejection rate as the proportion of replications for which the p-value (comparing the causal model to the non-causal sharing model) was less than 0.05.

**Table 1:**
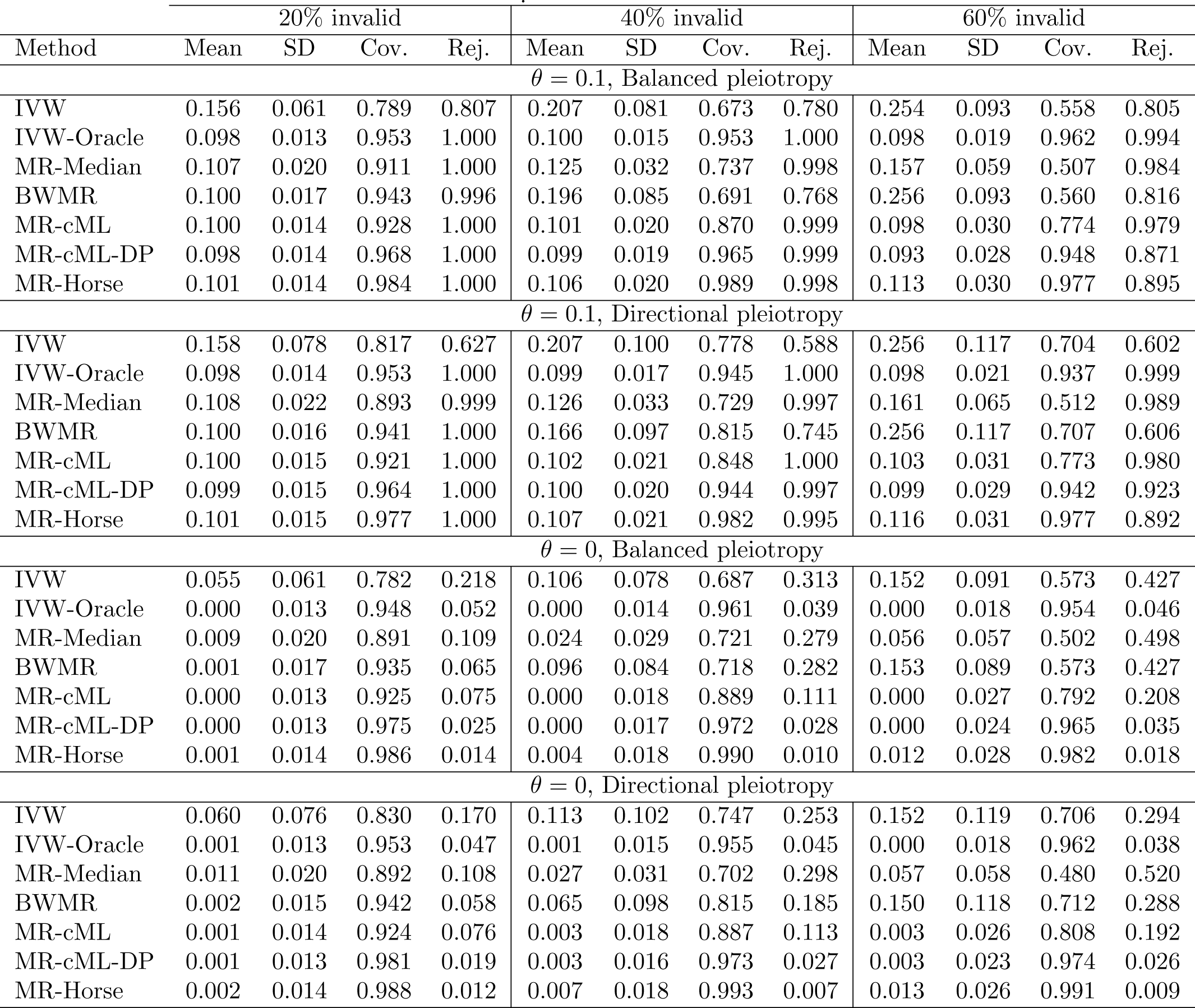
Results of simulations for the scenarios with 100 genetic instruments. Reported is the mean estimate (Mean), standard deviation of estimates (SD), proportion of replications that the 95% confidence interval (or credible interval in the case of MR-Horse) contained the true causal effect (Cov.) and the proportion of replications that the 95% confidence interval (or credible interval) did not contain zero (Rej.)

**Table 2:** Results of simulations for the scenarios with 20 genetic instruments. Reported is the mean estimate (Mean), standard deviation of estimates (SD), proportion of replications that the 95% confidence interval (or credible interval in the case of MR-Horse) contained the true causal effect (Cov.) and the proportion of replications that the 95% confidence interval (or credible interval) did not contain zero (Rej.)

| Method | 20% invalid |  |  |  | 40% invalid |  |  |  | 60% invalid |  |  |  |
| --- | --- | --- | --- | --- | --- | --- | --- | --- | --- | --- | --- | --- |
|  | Mean | SD | Cov. | Rej. | Mean | SD | Cov. | Rej. | Mean | SD | Cov. | Rej. |
| $\theta = 0.1$, Balanced pleiotropy | | | | | | | | | | | | |
| IVW | 0.150 | 0.135 | 0.897 | 0.307 | 0.190 | 0.178 | 0.869 | 0.262 | 0.244 | 0.209 | 0.853 | 0.277 |
| IVW-Oracle | 0.100 | 0.029 | 0.965 | 0.899 | 0.100 | 0.034 | 0.959 | 0.819 | 0.099 | 0.041 | 0.970 | 0.610 |
| MR-Median | 0.109 | 0.048 | 0.919 | 0.750 | 0.129 | 0.086 | 0.806 | 0.716 | 0.198 | 0.198 | 0.631 | 0.730 |
| BWMR | 0.135 | 0.120 | 0.895 | 0.456 | 0.188 | 0.177 | 0.870 | 0.263 | 0.248 | 0.207 | 0.847 | 0.277 |
| MR-cML | 0.101 | 0.033 | 0.919 | 0.905 | 0.100 | 0.042 | 0.874 | 0.807 | 0.096 | 0.061 | 0.802 | 0.632 |
| MR-cML-DP | 0.097 | 0.032 | 0.951 | 0.806 | 0.093 | 0.040 | 0.935 | 0.600 | 0.085 | 0.053 | 0.914 | 0.377 |
| MR-Horse | 0.102 | 0.034 | 0.991 | 0.690 | 0.106 | 0.044 | 0.992 | 0.381 | 0.122 | 0.082 | 0.989 | 0.195 |
| $\theta = 0.1$, Directional pleiotropy | | | | | | | | | | | | |
| IVW | 0.157 | 0.166 | 0.902 | 0.265 | 0.199 | 0.221 | 0.911 | 0.188 | 0.261 | 0.268 | 0.865 | 0.230 |
| IVW-Oracle | 0.099 | 0.030 | 0.957 | 0.891 | 0.099 | 0.035 | 0.956 | 0.763 | 0.097 | 0.042 | 0.970 | 0.568 |
| MR-Median | 0.109 | 0.053 | 0.893 | 0.749 | 0.132 | 0.093 | 0.808 | 0.712 | 0.201 | 0.204 | 0.599 | 0.713 |
| BWMR | 0.134 | 0.128 | 0.896 | 0.528 | 0.195 | 0.216 | 0.885 | 0.213 | 0.262 | 0.265 | 0.875 | 0.220 |
| MR-cML | 0.101 | 0.033 | 0.924 | 0.909 | 0.101 | 0.042 | 0.876 | 0.780 | 0.098 | 0.059 | 0.817 | 0.628 |
| MR-cML-DP | 0.097 | 0.033 | 0.957 | 0.830 | 0.095 | 0.041 | 0.934 | 0.608 | 0.089 | 0.053 | 0.911 | 0.401 |
| MR-Horse | 0.102 | 0.034 | 0.985 | 0.722 | 0.107 | 0.045 | 0.993 | 0.351 | 0.123 | 0.077 | 0.994 | 0.170 |
| $\theta = 0$, Balanced pleiotropy | | | | | | | | | | | | |
| IVW | 0.046 | 0.132 | 0.894 | 0.106 | 0.100 | 0.176 | 0.882 | 0.118 | 0.147 | 0.204 | 0.836 | 0.164 |
| IVW-Oracle | -0.001 | 0.027 | 0.969 | 0.031 | 0.000 | 0.032 | 0.959 | 0.041 | 0.000 | 0.040 | 0.965 | 0.035 |
| MR-Median | 0.009 | 0.044 | 0.915 | 0.085 | 0.033 | 0.093 | 0.791 | 0.209 | 0.093 | 0.181 | 0.587 | 0.413 |
| BWMR | 0.032 | 0.118 | 0.888 | 0.112 | 0.097 | 0.176 | 0.877 | 0.123 | 0.147 | 0.204 | 0.843 | 0.157 |
| MR-cML | -0.001 | 0.029 | 0.939 | 0.061 | 0.000 | 0.039 | 0.890 | 0.110 | 0.000 | 0.055 | 0.835 | 0.165 |
| MR-cML-DP | -0.001 | 0.027 | 0.975 | 0.025 | 0.000 | 0.035 | 0.955 | 0.045 | 0.000 | 0.047 | 0.942 | 0.058 |
| MR-Horse | 0.000 | 0.030 | 0.986 | 0.014 | 0.005 | 0.041 | 0.996 | 0.004 | 0.017 | 0.070 | 0.986 | 0.014 |
| $\theta = 0$, Directional pleiotropy | | | | | | | | | | | | |
| IVW | 0.047 | 0.174 | 0.883 | 0.117 | 0.096 | 0.227 | 0.888 | 0.112 | 0.153 | 0.268 | 0.871 | 0.129 |
| IVW-Oracle | -0.001 | 0.028 | 0.957 | 0.043 | 0.001 | 0.031 | 0.961 | 0.039 | 0.000 | 0.040 | 0.963 | 0.037 |
| MR-Median | 0.008 | 0.048 | 0.907 | 0.093 | 0.031 | 0.087 | 0.813 | 0.187 | 0.097 | 0.216 | 0.591 | 0.409 |
| BWMR | 0.025 | 0.131 | 0.886 | 0.114 | 0.088 | 0.219 | 0.884 | 0.116 | 0.152 | 0.265 | 0.877 | 0.123 |
| MR-cML | 0.000 | 0.030 | 0.932 | 0.068 | 0.002 | 0.037 | 0.901 | 0.099 | 0.003 | 0.055 | 0.827 | 0.173 |
| MR-cML-DP | 0.000 | 0.028 | 0.969 | 0.031 | 0.001 | 0.034 | 0.958 | 0.042 | 0.002 | 0.047 | 0.938 | 0.062 |
| MR-Horse | 0.001 | 0.031 | 0.992 | 0.008 | 0.006 | 0.040 | 0.995 | 0.005 | 0.019 | 0.072 | 0.995 | 0.005 |

**Table 3:** Results of simulations for the genome-wide simulation scenarios with either *J* = 100 or *J* = 20 genetic variants associated with the exposure. Reported is the mean estimate (Mean), standard deviation of estimates (SD), proportion of replications that the 95% confidence interval (or credible interval in the cases of CAUSE and MR-Horse) contained the true causal effect (Cov.) and the proportion of replications that the 95% confidence interval (or credible interval) did not contain zero or, in the case of CAUSE, the proportion of replications that the causal null hypothesis was rejected (Rej.)

| Method | 20% invalid |  |  |  | 40% invalid |  |  |  | 60% invalid |  |  |  |
| --- | --- | --- | --- | --- | --- | --- | --- | --- | --- | --- | --- | --- |
|  | Mean | SD | Cov. | Rej. | Mean | SD | Cov. | Rej. | Mean | SD | Cov. | Rej. |
| $J = 100, \theta = 0.1$ | | | | | | | | | | | | |
| IVW | 0.168 | 0.047 | 0.700 | 0.960 | 0.227 | 0.065 | 0.465 | 0.955 | 0.301 | 0.084 | 0.260 | 0.985 |
| MR-Median | 0.107 | 0.014 | 0.925 | 1.000 | 0.119 | 0.027 | 0.760 | 1.000 | 0.168 | 0.071 | 0.410 | 1.000 |
| BWMR | 0.111 | 0.029 | 0.935 | 1.000 | 0.195 | 0.075 | 0.575 | 0.950 | 0.298 | 0.085 | 0.270 | 0.990 |
| MR-cML | 0.100 | 0.010 | 0.960 | 1.000 | 0.100 | 0.013 | 0.895 | 1.000 | 0.102 | 0.022 | 0.810 | 0.990 |
| MR-cML-DP | 0.099 | 0.009 | 0.975 | 1.000 | 0.099 | 0.013 | 0.960 | 1.000 | 0.099 | 0.021 | 0.930 | 0.970 |
| CAUSE | 0.101 | 0.010 | 0.990 | 0.920 | 0.103 | 0.014 | 0.980 | 0.885 | 0.111 | 0.025 | 0.960 | 0.760 |
| MR-Horse | 0.102 | 0.010 | 0.995 | 1.000 | 0.107 | 0.014 | 0.980 | 1.000 | 0.123 | 0.023 | 0.960 | 0.990 |
| $J = 100, \theta = 0$ | | | | | | | | | | | | |
| IVW | 0.069 | 0.047 | 0.695 | 0.305 | 0.128 | 0.066 | 0.465 | 0.535 | 0.202 | 0.084 | 0.265 | 0.735 |
| MR-Median | 0.008 | 0.014 | 0.925 | 0.075 | 0.020 | 0.027 | 0.760 | 0.240 | 0.068 | 0.069 | 0.410 | 0.590 |
| BWMR | 0.011 | 0.029 | 0.940 | 0.060 | 0.095 | 0.075 | 0.575 | 0.425 | 0.198 | 0.085 | 0.280 | 0.720 |
| MR-cML | 0.000 | 0.009 | 0.965 | 0.035 | 0.001 | 0.013 | 0.915 | 0.085 | 0.003 | 0.020 | 0.810 | 0.190 |
| MR-cML-DP | 0.000 | 0.009 | 0.980 | 0.020 | 0.001 | 0.012 | 0.970 | 0.030 | 0.003 | 0.018 | 0.950 | 0.050 |
| CAUSE | 0.001 | 0.009 | 0.985 | 0.455 | 0.002 | 0.011 | 0.990 | 0.435 | 0.005 | 0.018 | 0.960 | 0.335 |
| MR-Horse | 0.002 | 0.010 | 0.995 | 0.005 | 0.008 | 0.014 | 0.985 | 0.015 | 0.023 | 0.023 | 0.955 | 0.045 |
| $J = 20, \theta = 0.1$ | | | | | | | | | | | | |
| IVW | 0.168 | 0.117 | 0.885 | 0.555 | 0.251 | 0.163 | 0.825 | 0.455 | 0.307 | 0.181 | 0.740 | 0.525 |
| MR-Median | 0.110 | 0.050 | 0.930 | 1.000 | 0.144 | 0.120 | 0.795 | 1.000 | 0.207 | 0.177 | 0.530 | 0.970 |
| BWMR | 0.137 | 0.099 | 0.905 | 0.770 | 0.228 | 0.168 | 0.835 | 0.595 | 0.298 | 0.176 | 0.735 | 0.540 |
| MR-cML | 0.099 | 0.011 | 0.955 | 1.000 | 0.100 | 0.019 | 0.885 | 0.990 | 0.094 | 0.034 | 0.840 | 0.960 |
| MR-cML-DP | 0.098 | 0.011 | 0.980 | 0.995 | 0.098 | 0.018 | 0.920 | 0.960 | 0.090 | 0.028 | 0.890 | 0.905 |
| CAUSE | 0.097 | 0.014 | 0.995 | 0.725 | 0.105 | 0.042 | 0.990 | 0.555 | 0.113 | 0.074 | 1.000 | 0.335 |
| MR-Horse | 0.101 | 0.015 | 0.990 | 0.980 | 0.109 | 0.030 | 1.000 | 0.925 | 0.138 | 0.111 | 0.975 | 0.745 |
| $J = 20, \theta = 0$ | | | | | | | | | | | | |
| IVW | 0.068 | 0.117 | 0.885 | 0.115 | 0.151 | 0.164 | 0.825 | 0.175 | 0.207 | 0.182 | 0.740 | 0.260 |
| MR-Median | 0.010 | 0.050 | 0.930 | 0.070 | 0.044 | 0.121 | 0.800 | 0.200 | 0.107 | 0.178 | 0.540 | 0.460 |
| BWMR | 0.037 | 0.099 | 0.900 | 0.100 | 0.128 | 0.169 | 0.835 | 0.165 | 0.198 | 0.177 | 0.735 | 0.265 |
| MR-cML | 0.000 | 0.010 | 0.955 | 0.045 | 0.002 | 0.014 | 0.895 | 0.105 | -0.001 | 0.029 | 0.900 | 0.100 |
| MR-cML-DP | -0.001 | 0.010 | 0.980 | 0.020 | 0.002 | 0.013 | 0.930 | 0.070 | 0.000 | 0.018 | 0.955 | 0.045 |
| CAUSE | 0.000 | 0.010 | 0.990 | 0.395 | 0.002 | 0.014 | 0.965 | 0.320 | 0.010 | 0.065 | 0.990 | 0.210 |
| MR-Horse | 0.001 | 0.015 | 0.990 | 0.010 | 0.008 | 0.050 | 1.000 | 0.000 | 0.033 | 0.090 | 0.990 | 0.010 |

**Table 4:** Results of simulations for the multivariable scenarios (relating to the causal effect of *X*_1_ only). Reported is the mean estimate (Mean), standard deviation of estimates (SD), proportion of replications that the 95% confidence interval (or credible interval in the case of MVMR-Horse) contained the true causal effect (Cov.) and the proportion of replications that the 95% confidence interval (or credible interval) did not contain zero (Rej.)

| Method | 20% invalid |  |  |  | 40% invalid |  |  |  | 60% invalid |  |  |  |
| --- | --- | --- | --- | --- | --- | --- | --- | --- | --- | --- | --- | --- |
|  | Mean | SD | Cov. | Rej. | Mean | SD | Cov. | Rej. | Mean | SD | Cov. | Rej. |
| $\theta_1 = 0.1$, Balanced pleiotropy | | | | | | | | | | | | |
| IVW | 0.155 | 0.066 | 0.785 | 0.805 | 0.204 | 0.078 | 0.720 | 0.765 | 0.229 | 0.088 | 0.670 | 0.700 |
| IVW-Oracle | 0.100 | 0.014 | 0.950 | 1.000 | 0.098 | 0.016 | 0.965 | 1.000 | 0.097 | 0.020 | 0.970 | 0.990 |
| MVMR-Median | 0.107 | 0.020 | 0.935 | 1.000 | 0.117 | 0.027 | 0.845 | 1.000 | 0.137 | 0.047 | 0.700 | 0.975 |
| GRAPPLE | 0.109 | 0.032 | 0.990 | 0.775 | 0.169 | 0.068 | 0.875 | 0.685 | 0.213 | 0.088 | 0.795 | 0.660 |
| MVMR-cML-DP | 0.101 | 0.015 | 0.975 | 1.000 | 0.099 | 0.020 | 0.945 | 1.000 | 0.096 | 0.031 | 0.935 | 0.850 |
| MVMR-Horse | 0.102 | 0.015 | 0.980 | 1.000 | 0.105 | 0.020 | 1.000 | 0.995 | 0.114 | 0.032 | 0.980 | 0.880 |
| $\theta_1 = 0.1$, Directional pleiotropy | | | | | | | | | | | | |
| IVW | 0.154 | 0.077 | 0.855 | 0.620 | 0.208 | 0.109 | 0.790 | 0.585 | 0.229 | 0.105 | 0.805 | 0.535 |
| IVW-Oracle | 0.098 | 0.014 | 0.955 | 1.000 | 0.100 | 0.017 | 0.965 | 1.000 | 0.098 | 0.022 | 0.940 | 0.990 |
| MVMR-Median | 0.105 | 0.019 | 0.970 | 1.000 | 0.119 | 0.032 | 0.820 | 0.995 | 0.142 | 0.051 | 0.670 | 0.965 |
| GRAPPLE | 0.109 | 0.031 | 0.995 | 0.680 | 0.175 | 0.087 | 0.900 | 0.510 | 0.217 | 0.103 | 0.830 | 0.490 |
| MVMR-cML-DP | 0.100 | 0.016 | 0.975 | 1.000 | 0.103 | 0.020 | 0.955 | 0.990 | 0.106 | 0.030 | 0.950 | 0.935 |
| MVMR-Horse | 0.101 | 0.015 | 0.975 | 1.000 | 0.109 | 0.021 | 0.990 | 0.995 | 0.121 | 0.034 | 0.970 | 0.865 |
| $\theta_1 = 0$, Balanced pleiotropy | | | | | | | | | | | | |
| IVW | 0.055 | 0.061 | 0.770 | 0.230 | 0.103 | 0.078 | 0.690 | 0.310 | 0.138 | 0.086 | 0.705 | 0.295 |
| IVW-Oracle | -0.001 | 0.013 | 0.975 | 0.025 | 0.003 | 0.017 | 0.925 | 0.075 | -0.004 | 0.020 | 0.950 | 0.050 |
| MVMR-Median | 0.004 | 0.017 | 0.960 | 0.040 | 0.020 | 0.028 | 0.775 | 0.225 | 0.030 | 0.042 | 0.750 | 0.250 |
| GRAPPLE | 0.006 | 0.027 | 1.000 | 0.000 | 0.070 | 0.068 | 0.835 | 0.165 | 0.118 | 0.083 | 0.775 | 0.225 |
| MVMR-cML-DP | -0.001 | 0.014 | 0.970 | 0.030 | 0.002 | 0.020 | 0.955 | 0.045 | -0.008 | 0.028 | 0.930 | 0.070 |
| MVMR-Horse | 0.000 | 0.014 | 1.000 | 0.000 | 0.008 | 0.022 | 0.990 | 0.010 | 0.007 | 0.030 | 0.980 | 0.020 |
| $\theta_1 = 0$, Directional pleiotropy | | | | | | | | | | | | |
| IVW | 0.049 | 0.072 | 0.870 | 0.130 | 0.096 | 0.104 | 0.795 | 0.205 | 0.133 | 0.115 | 0.810 | 0.190 |
| IVW-Oracle | 0.000 | 0.014 | 0.940 | 0.060 | -0.002 | 0.017 | 0.960 | 0.040 | -0.001 | 0.021 | 0.960 | 0.040 |
| MVMR-Median | 0.006 | 0.020 | 0.945 | 0.055 | 0.014 | 0.030 | 0.830 | 0.170 | 0.031 | 0.041 | 0.770 | 0.230 |
| GRAPPLE | 0.011 | 0.028 | 1.000 | 0.000 | 0.059 | 0.083 | 0.915 | 0.085 | 0.112 | 0.111 | 0.840 | 0.160 |
| MVMR-cML-DP | 0.001 | 0.016 | 0.950 | 0.050 | 0.000 | 0.020 | 0.955 | 0.045 | 0.002 | 0.031 | 0.910 | 0.090 |
| MVMR-Horse | 0.002 | 0.016 | 0.970 | 0.030 | 0.005 | 0.021 | 0.995 | 0.005 | 0.013 | 0.031 | 0.970 | 0.030 |

In the primary univariable simulations, the mean estimates from MR-cML, MR-cML-DP and MR-Horse were close to IVW-Oracle, whereas IVW and BWMR showed considerable bias from 40% pleiotropy and MR-Median at 60% pleiotropy. Although MR-cML retained nominal coverage and type I error rates up to 40% pleiotropy, these were considerably away from nominal levels at 60% pleiotropy. MR-cML-DP retained nominal coverage and type I error rates for most *J* = 100 scenarios, however had undercoverage and inflated type I error rates in the *J* = 20 scenarios at the higher pleiotropy levels. In comparison, MR-Horse retained coverage over the nominal level, and type I error rates below the nominal level, across all scenarios. In general, MR-cML-DP showed slightly higher power, whereas MR-Horse showed slightly higher coverage, particularly in the *J* = 20 scenarios.

In the genome-wide simulation study, MR-Horse had comparable power and bias to CAUSE, with substantially lower type I error rates. It again had slightly higher coverage and lower type I error rates compared with MR-cML-DP, and outperformed all other methods across each metric.

In the multivariable simulation study, MVMR-Horse outperformed IVW, MVMR-Median and GRAPPLE in terms of bias, precision and type I error rates in all scenarios. Similar to the univariable settings, MVMR-Horse retained type I error rates below the nominal level in all scenarios, with the trade-off of lower power compared with MVMR-cML-DP. MVMR-cML-DP had inflated type I error rates in the higher pleiotropy scenarios.

### Applied example results

Figure 2 shows the estimated log odds ratio of each outcome per standard deviation increase in the genetically-predicted levels of each exposure, as well as the 95% confidence intervals (IVW, MR-cML, MR-cML-DP) or 95% credible intervals (CAUSE, MR-Horse). For the exposure-outcome combinations with established causality, the results from each method were broadly in line across the different methods. For LDL-T2D, CAUSE did not show evidence of a causal effect. MR-Horse agreed with MR-cML(-DP) that increased LDL may reduce T2D risk, although with a 95% credible interval that came closer to the null. The other conflicting result was that CAUSE and MR-cML showed evidence of a causal effect of TG on T2D risk. MR-cML-DP and MR-Horse produced a null result for TG-T2D, suggesting that CAUSE and MR-cML may be producing a false positive result in this case. All methods showed a causal effect of BW on the risk of CAD and T2D, although with the estimates from CAUSE much closer to the null than the others. Finally, all methods gave a null result for the causal effect of LDL, TG and BW on the risk of asthma. However, MR-Horse showed a possible causal relationship between fasting glucose and risk of asthma, agreeing with MR-cML(-DP).

**Figure 2:**
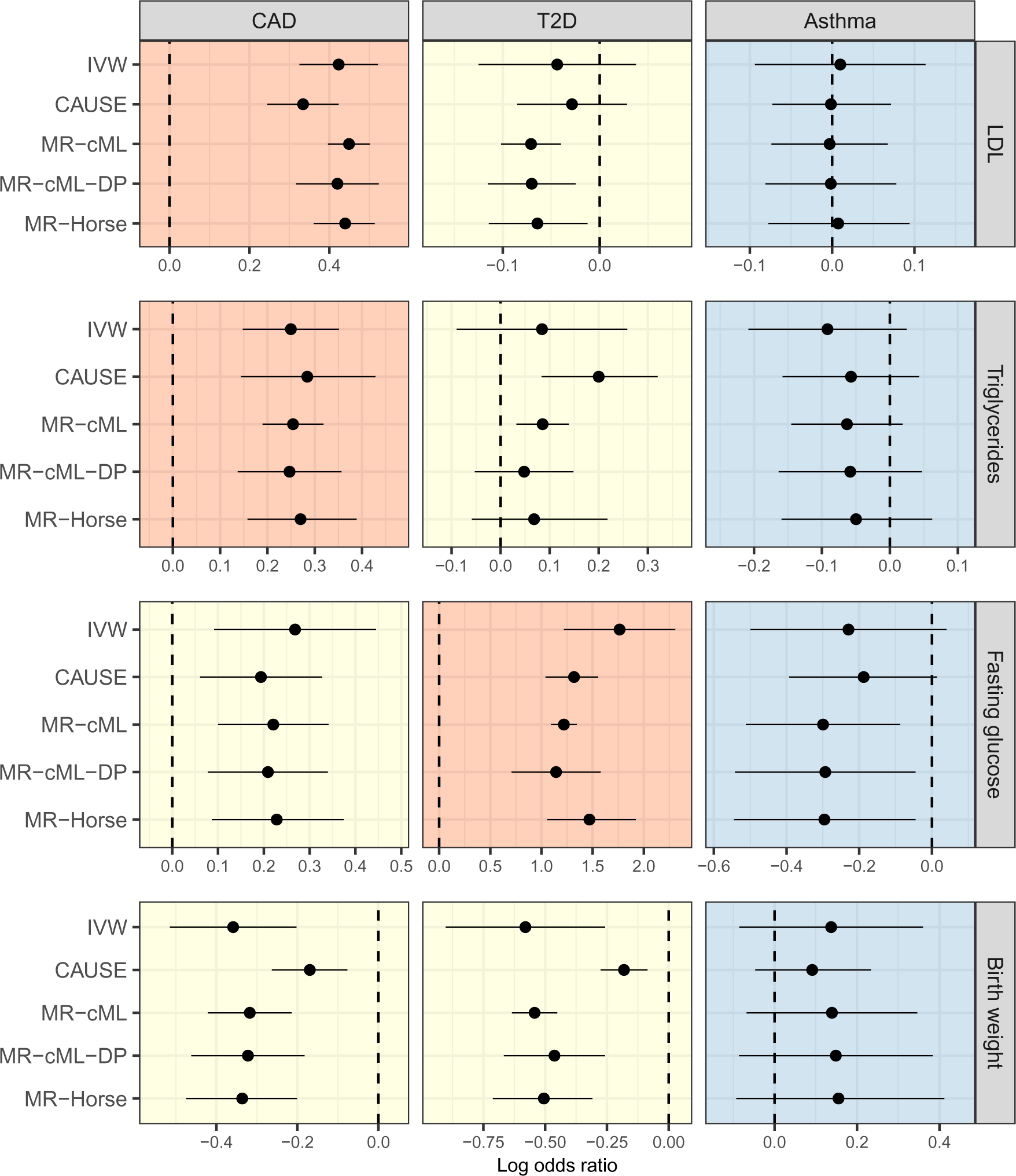
Mendelian randomization estimates and 95% confidence intervals (IVW, MR-cML, MR-cML-DP) or 95% credible intervals (CAUSE, MR-Horse) for the different exposure-outcome combinations. Plots coloured in red indicate exposure-outcome relationships considered causal. Plots coloured in yellow indicate exposure-outcome pairs with unknown causal relationships. Plots coloured in blue indicate exposure-outcome relationships considered not causal. In each plot, the vertical dashed line indicates a log odds ratio of zero.

## Discussion

In this paper, we have presented a Bayesian framework for performing Mendelian randomization with summary level data which is robust to general forms of instrument invalidity. The approach does not require the InSIDE assumption and so it can handle correlated pleiotropy. The method is easily generalised to the multivariable Mendelian randomization setting, providing another robust method for multivariable studies. Our simulation studies showed that the approach has favorable performance in comparison with commonly used methods across a variety of settings.

Our approach has similarities to previously proposed Bayesian approaches, but has been designed to avoid many of the restrictions that these methods pose. Berzuini et al. (2020) first proposed a Bayesian framework for Mendelian randomization with pleiotropy and suggested the use of the horseshoe prior to account for the presence of some valid and some invalid instruments. However, their method requires individual level data and makes the InSIDE assumption. Zhao et al. (2019) and Bucur et al. (2020) developed versions of the Berzuini et al. approach using summary level data, again requiring InSIDE to be satisfied. Cheng et al. (2021) introduced a correlation parameter into a Bayesian model to account for correlated pleiotropy, however their model requires that pleiotropic effects are either all correlated or all uncorrelated with the variant-exposure effects. The methods of Morrison et al. (2020) and Cheng et al. (2022) are able to account for both correlated and uncorrelated pleiotropy, but make a further assumption that the genetic instruments are associated with the exposure either entirely through the confounders, or entirely through a direct effect. Our approach utilises the horseshoe prior as suggested by Berzuini et al., can be applied using summary level data, and can account for both correlated and uncorrelated pleiotropy without any assumption on the pathways between the genetic instruments and exposure. Furthermore, we have shown how to extend the framework to be used for multivariable Mendelian randomization.

In comparison to frequentist approaches for Mendelian randomization, our method is based on the same causal model as that used by Xue et al. (2021), which we believe is a very general model for instrument invalidity. The performance of our method in simulation studies was comparable to Xue et al.’s method when the latter was applied with the data perturbation extension. However, particularly in high pleiotropy settings, our Bayesian approach retained lower type I error rates which were uniformly below the nominal level, with the trade-off being lower power than the frequentist approach. The increased conservatism in the Bayesian approach supports the assertion that the horseshoe prior is able to avoid shrinkage of the pleiotropic effects pertaining to invalid instruments. Besides improved type I error control, another advantage of our framework is that it allows for the model specification to be adapted to specific settings. For example, the prior distribution of the global shrinkage parameter could be adjusted, or set at a fixed value, based on prior knowledge of the level and nature of pleiotropy. As long as these prior assumptions are reasonable, such incorporation of prior knowledge would be expected to reduce the uncertainty in the estimates and increase power. Although the implementation of the method in this paper has used mostly weakly-informative priors for the model parameters, an interesting area for future development of the approach would be to explore the use of domain knowledge to inform the model inputs.

Not taken into account in our model are sample overlap and linkage disequilibrium. Most of the methods which are commonly used for Mendelian randomization assume independent genetic variants, and achieve this by pruning candidate instruments according to an *r*^2^ threshold. In general, the use of correlated variants is unlikely to add substantially more information to an analysis, and methods which are able to incorporate linkage disequilibrium tend to be very sensitive to mis-specification of genetic correlations (Burgess et al., 2017). Nonetheless, there are situations where correlated variants may be desirable, such as in a *cis*-Mendelian randomization study exploring drug targets (Schmidt et al., 2020). In this case, correlation estimates could be incorporated into our model by specifying a joint distribution for the genetic variant-exposure association estimates and for the genetic variant-outcome association estimates which included genetic correlations, as is done by Cheng et al. (2022) to account for linkage disequilibrium.

Sample overlap is more likely to be an issue in practice, since many published GWAS results are from meta-analyses which include common data sources such as the UK Biobank (Sudlow et al., 2015). The models of Morrison et al. (2020) and Cheng et al. (2022) account for sample overlap by including correlation between 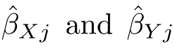 in a joint distribution. Our model could be adapted in a similar way to account for sample overlap if it was believed to be substantial.

Assumptions that we have made are of the linearity of the effect of the exposures on the outcome, and homogeneity or no effect modification. Furthermore, an implicit assumption that we have made in developing the method and simulation study is that all traits are continuous. In practice, summary level Mendelian randomization methods can be performed with categorical traits, for example by taking genetic association estimates from logistic regression. This may bias causal effect estimates because of the non-collapsibility of odds ratios, however the bias will tend to be toward the null, at least in the univariable case (Bowden et al., 2017). One further drawback to Bayesian methods generally is that they tend to be computationally intensive. However, it should be noted that MR-Horse was quicker to run than both MR-cML-DP and CAUSE.

Overall, our Bayesian framework is an effective and flexible method for both univariable and multivariable Mendelian randomization. It provides an important sensitivity analysis to these studies which is robust to violations of the InSIDE assumption and directional pleiotropy. Furthermore, it can be easily adapted to handle sample overlap or linkage disequilibrium.

## Acknowledgments

SB is supported by the Wellcome Trust (225790/Z/22/Z) and the United Kingdom Research and Innovation Medical Research Council (MC UU 00002/7). This research was supported by the National Institute for Health Research Cambridge Biomedical Research Centre (NIHR203312). The views expressed are those of the authors and not necessarily those of the National Institute for Health Research or the Department of Health and Social Care.

The authors acknowledge the Sydney Informatics Hub and the University of Sydney’s high per-formance computing cluster, Artemis, for providing the computing resources that have contributed to the results reported herein.

## Data and code availability

All the data used in this paper are publicly available and can be accessed via the references given. R code for performing the proposed methods, and for reproducing the simulation results and applied analyses, can be found at https://github.aj-grant/mrhorse.

